# Spatial Organization of Rho GTPase signaling by RhoGEF/RhoGAP proteins

**DOI:** 10.1101/354316

**Authors:** Paul M. Müller, Juliane Rademacher, Richard D. Bagshaw, Keziban M. Alp, Girolamo Giudice, Louise E. Heinrich, Carolin Barth, Rebecca L. Eccles, Marta Sanchez-Castro, Lennart Brandenburg, Geraldine Mbamalu, Monika Tucholska, Lisa Spatt, Celina Wortmann, Maciej T. Czajkowski, Robert-William Welke, Sunqu Zhang, Vivian Nguyen, Trendelina Rrustemi, Philipp Trnka, Kiara Freitag, Brett Larsen, Oliver Popp, Philipp Mertins, Chris Bakal, Anne-Claude Gingras, Olivier Pertz, Frederick P. Roth, Karen Colwill, Tony Pawson, Evangelia Petsalaki, Oliver Rocks

## Abstract

Rho GTPases control cell morphogenesis and thus fundamental processes in all eukaryotes. They are regulated by 145 RhoGEF and RhoGAP multi-domain proteins in humans. How the Rho signaling system is organized to generate localized responses in cells and prevent their spreading is not understood. Here, we systematically characterized the substrate specificities, localization and interactome of the RhoGEFs/RhoGAPs and revealed their critical role in contextualizing and spatially delimiting Rho signaling. They localize to multiple compartments providing positional information, are extensively interconnected to jointly coordinate their signaling networks and are widely autoinhibited to remain sensitive to local activation. RhoGAPs exhibit lower substrate specificity than RhoGEFs and may contribute to preserving Rho activity gradients. Our approach led us to uncover a multi-RhoGEF complex downstream of G-protein-coupled receptors controlling a Cdc42/RhoA crosstalk. The spatial organization of Rho signaling thus differs from other small GTPases and expands the repertoire of mechanisms governing localized signaling activity.

---

Rho GTPases coordinate changes in cytoskeletal architecture to regulate cell shape. They drive fundamental aspects of cell behavior in all eukaryotes, including motility, cytokinesis and tissue morphogenesis(*1, 2*). Defects in Rho signaling have been widely found in cancer metastasis and other serious diseases(*3*). Rho proteins typically cycle between an inactive GDP-bound and an active GTP-bound form(*4*). Upon activation, they bind effector proteins to elicit cytoskeletal remodeling. Their activity cycle is initiated by guanine nucleotide exchange factors (RhoGEFs)(*5, 6*) and terminated by GTPase activating proteins (RhoGAPs)(*7*). In addition, guanine nucleotide dissociation inhibitors (RhoGDIs) sequester the GTPases in the cytosol to render them inactive(*8*). With 145 members, the RhoGEF/RhoGAP multi-domain proteins by far outnumber the 12 classical Rho GTPase switch proteins they regulate, allowing for complex control of Rho signaling activity and specificity (Fig. S1).

Rho signaling responses in cells are highly localized. A critical aspect of Rho biology is therefore its spatiotemporal control(*9*). Rho activities have been observed in distinct subcellular zones(*10–13*), with several GTPases operating simultaneously. Cell morphogenesis thus involves the concerted action of multiple Rho family members and their regulators, which together form complex local networks(*14, 15*). However, our knowledge how the Rho signaling system is orchestrated to give rise to such spatially confined cell responses is limited and stems from studies on individual Rho regulators, while a systems-level view is lacking.

Here, we undertook a comprehensive characterization of the RhoGEFs and RhoGAPs, including their substrate specificities, subcellular localization and interactomes. Our study places the regulators in functional context and enabled us to uncover emergent spatial organization principles that contribute to our understanding how Rho activity zones are dynamically maintained in cells.

## RhoGAPs exhibit lower substrate selectivity than RhoGEFs in a family-wide activity screen

We generated an expression library comprising 141 mammalian full-length RhoGEF/RhoGAP cDNAs, almost all of which represent the longest isoform known to exist (Fig. S1; Table S1).

As a first step, we systematically characterized the substrate specificities of the regulators to link them to their downstream pathways. An extensive literature survey and curation of claimed specificities revealed an incomplete data landscape with a high degree of conflict between reports (Table S2, Supplementary Text), highlighting the need for a standardized comprehensive analysis. We therefore developed a screening-compatible automated live-cell imaging assay using the latest generation of FRET-based biosensors for the canonical GTPases RhoA, Rac1 and Cdc42(*16–18*) (Fig. S2C-H). This approach enables the analysis of full-length regulators in their native cellular environment. We found catalytic activities for 45 out of 75 RhoGEFs and for 48 out of 63 RhoGAPs tested. In addition, the dual GEF/GAP proteins ABR and BCR both exhibited GAP activity (Fig. 1, Fig. S2A,B, Table S2). The active regulators included proteins whose substrate specificity was exclusive to one target (35 RhoGEFs, 31 RhoGAPs) but also many that regulated multiple Rho GTPases (10 RhoGEFs, 19 RhoGAPs). Our results thus not only reveal extensive promiscuity among the regulators, but also that in the GDP/GTP reaction cycle the inactivating RhoGAPs are less selective than the activating RhoGEFs (p-value=0.02)(Table S2). Promiscuity has been predominantly reported in the co-regulation of Cdc42 and Rac1, presumably because they control related cytoskeletal processes such as the formation of actin-rich protrusions at the leading edge (*19, 20*). However, we found a similar number of regulators that control RhoA together with Cdc42 and/or Rac1 (18 and 17 vs. 16), and thus GTPase combinations that elicit more diverse downstream responses, including RhoA-dependent regulation of actomyosin contractility. While we see high overall agreement of identified substrate specificities with existing data (70%, Table S2), we describe 10 novel activities. PLEKHG4B, for instance, is a strong exclusive Cdc42 GEF (see also Fig. 5) and SYDE2 a Rac1-specific GAP. The screen also revealed discrepancies with literature. The previously proposed representative Cdc42-specific GAPs ARHGAP1, ARHGAP17 or ARHGAP31 showed either no activity towards this GTPase in our assay or rather inactivated Rac1 more efficiently. In fact, none of the GAPs tested exhibited exclusive substrate specificity for Cdc42, although 21 GAPs promiscuously regulated Cdc42. By contrast, we found exclusive Cdc42 activity for a total of 12 GEFs.

**Figure 1.**
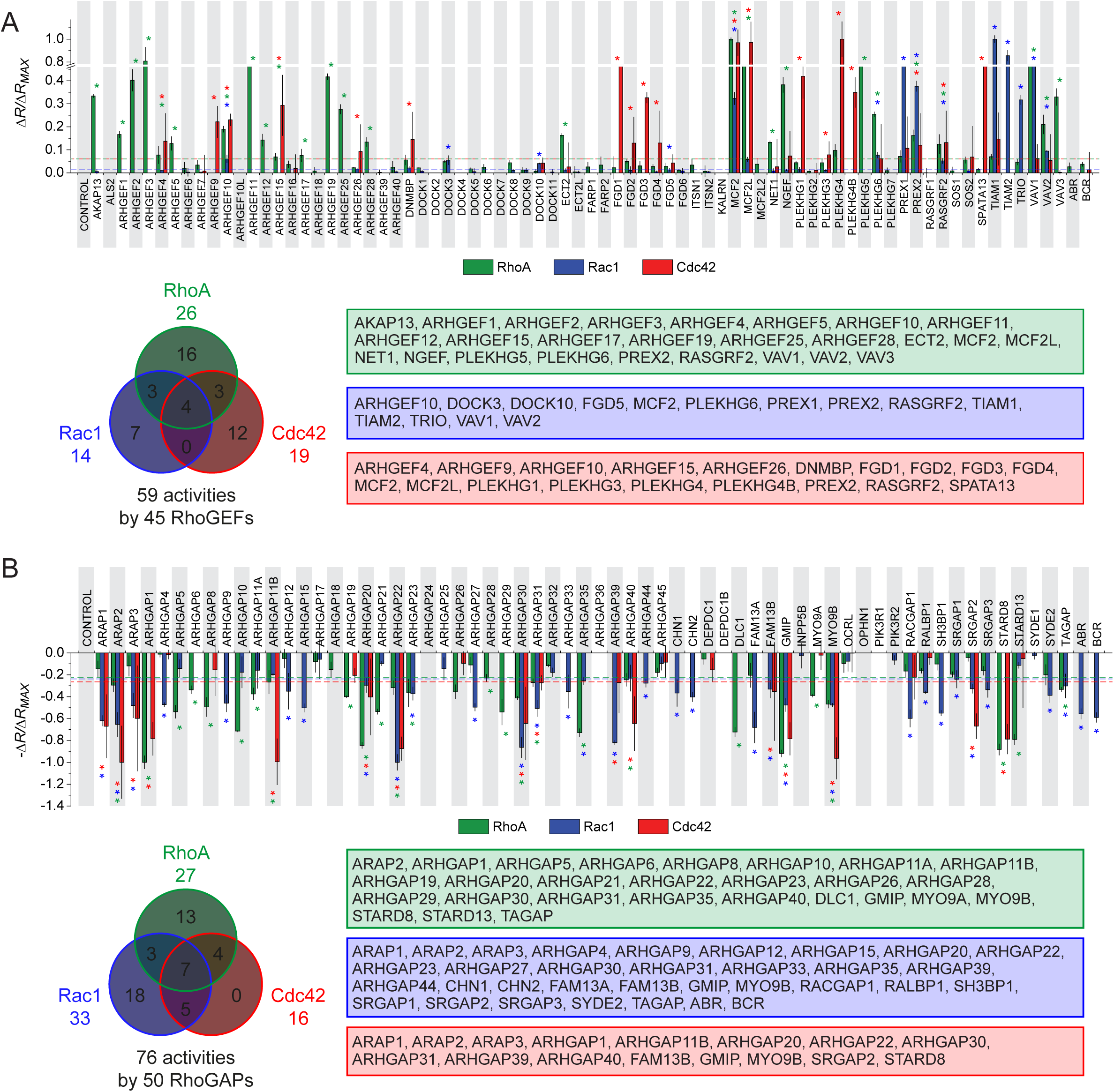
Family-wide RhoGEF and RhoGAP activity screens. (A) HEK293T cells were transfected with the mCherry-labeled RhoGEF cDNA library and the FRET sensors RhoA-2G, Rac1-2G, and Cdc42-2G together with RhoGDI. (B) RhoGDI shRNA-depleted HEK293T cells were transfected with the mCherry-labeled RhoGAP cDNA library and indicated FRET sensors. Values show changes in FRET ratio (δR) compared to control normalized to the maximal observed FRET ratio change (δR_MAX_). Mean ± SD (n=3) is shown throughout. Dashed lines indicate activity thresholds. Significance of values above (A) or below (B) the threshold was calculated by unpaired Student’s t-test versus control and Benjamini-Hochberg procedure, significant values are marked with an asterisk (*). For regulators with multiple activities, only those above 20% of the main activity were considered. Venn diagrams and lists of active regulators are given sorted by substrate GTPases: RhoA (green), Rac1 (blue), and Cdc42 (red).

## Autoinhibition is a common feature of RhoGEFs and RhoGAPs

Some RhoGEFs and RhoGAPs displayed minimal or no activity in our screen. A requirement for release from autoinhibition via mechanisms such as phosphorylation, protein or lipid interactions may account for the observed inactivity(*21*). To study whether autoinhibition is a general mechanism in controlling the Rho regulators we included shorter forms of 9 RhoGEFs and 10 RhoGAPs, which lack potential regulatory elements present in the longest isoforms. 13 of the shorter proteins indeed exhibited a higher catalytic efficiency than their longer counterparts (Fig. 2). ARHGAP9 normally only exhibits catalytic activity towards Rac1 as its longest isoform. Surprisingly, isoform 3, lacking an N-terminal SH3 domain, also inactivated Cdc42, suggesting that autoregulatory features might also affect the substrate selectivity of Rho regulators. Overall, our data shows that RhoGEFs and RhoGAPs are widely subject to autoinhibition.

**Figure 2.**
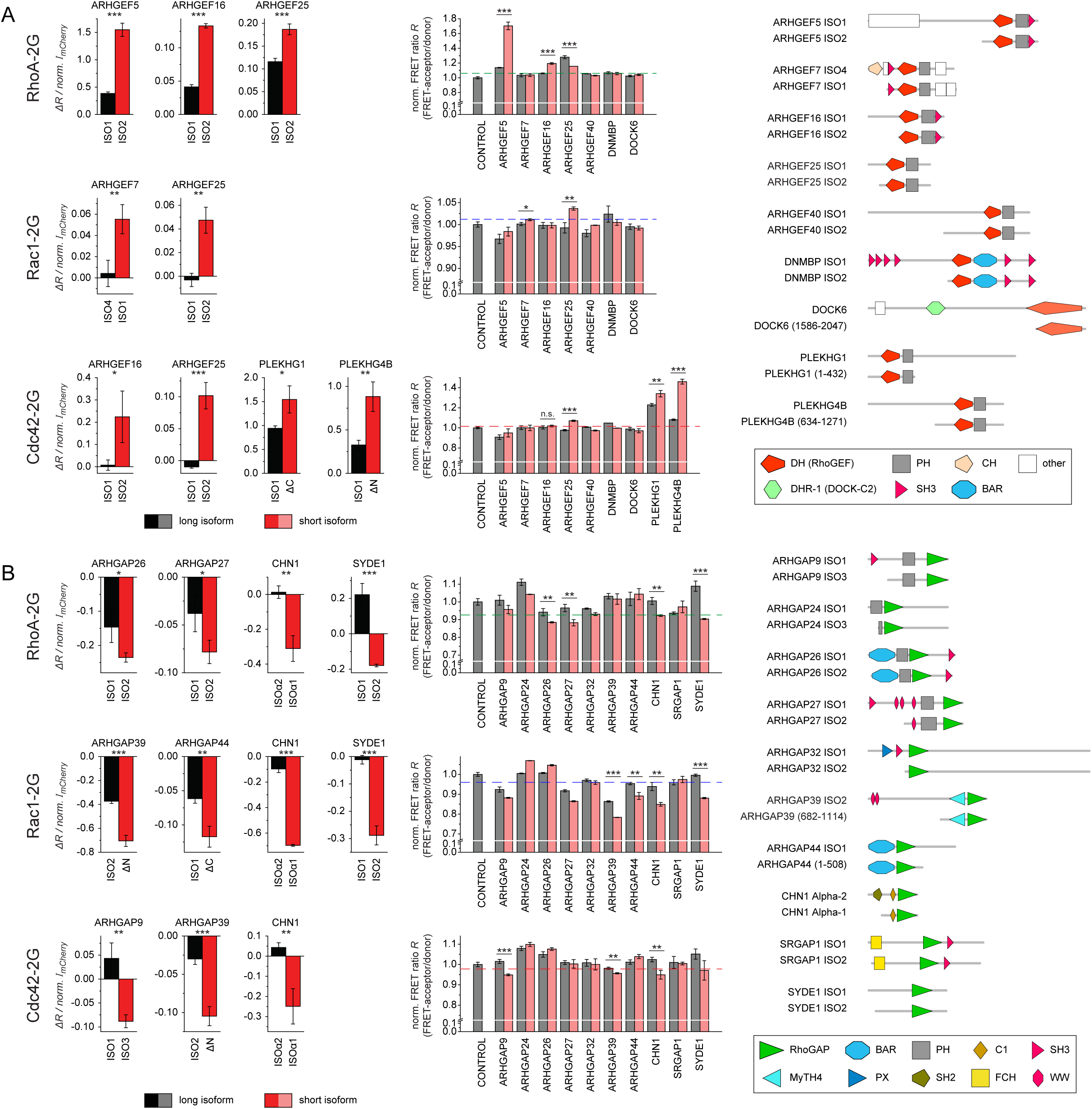
Autoinhibition is a common feature of RhoGEFs and RhoGAPs. Semiquantitative comparison of catalytic activities of full-length longest isoforms versus shorter isoforms or truncations of randomly selected 9 RhoGEFs (A) and 10 RhoGAPs (B). Experiments were performed as described in Figure 1 and Figure S2. Graphs on the left represent changes in FRET ratio (δR) normalized to RhoGEF or RhoGAP expression levels as determined by mCherry intensity. Graphs on the right show the FRET ratios normalized to control. Dashed lines indicate activity thresholds as in Figure 1. Values show mean ± SD (n=3; n=2 for ARHGEF40 and DNMBP}. Significance levels were calculated by unpaired Student’s t-tests as indicated: ***p<0.001, **p<0.01, *p<0.05, n.s.=not significant. Right panels: domain representations of the used constructs, as in Figure S1.

## RhoGEFs and RhoGAPs display diverse spatial distribution in cells

To understand how RhoGEFs and RhoGAPs provide spatial information to Rho signaling, we mapped the subcellular distribution of all 141 regulators in our library. We first used confocal live-cell microscopy to screen YFP fusion proteins in MDCK epithelial cell. This analysis revealed that over half of RhoGEFs and RhoGAPs (76/141) inherently localize to one or more distinct structures at steady-state, collectively decorating virtually all cellular compartments (Fig. 3A, Fig. S3, Table S3). For two proteins, ARHGAP9 and MCF2L2, we observed dependence of localization on the position of the fluorescent protein tag (Fig. S3). Interestingly, ARHGAP9-YFP localized to mitochondria while YFP-ARHGAP9 was enriched at cell junctions, perhaps because the N-terminal YFP fusion disrupted a mitochondrial targeting signal predicted for ARHGAP9 residues 1-14 (PredSL tool(*22*)). Indeed, ARHGAP9-YFP isoform 3, lacking residues 1-181, did not localize to mitochondria. We identified eight other cases in which different isoforms exhibited distinct localization patterns (Fig. S3).

**Figure 3.**
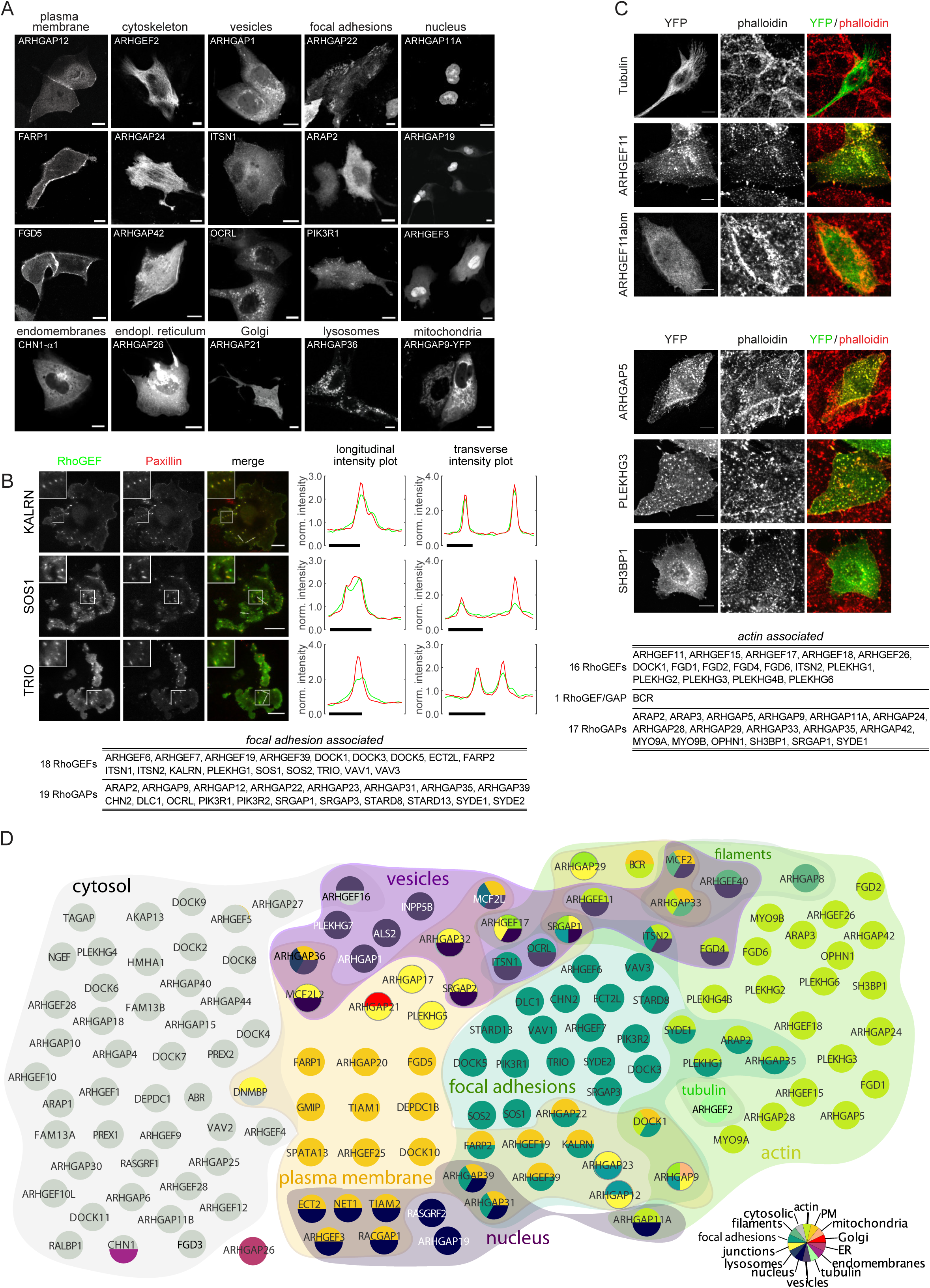
RhoGEFs and RhoGAPs display diverse spatial distribution in cells. Selection of images from confocal live-cell microscopy screen of YFP-tagged RhoGEF and RhoGAP subcellular localization (A), TIRF microscopy focal adhesion localization screen (B), and Cytochalasin D-treated actin colocacalization screen (C). (B) Image insets show higher magnifications of the boxed areas. Right panels: normalized fluorescence intensity profiles of longitudinal and transverse sections through focal adhesions, indicated by white lines in the merged images. (C) abm = actin binding mutant. Scale bars: (A) 10 µm, (B) white scale bars: 20 µm, black scale bars: 5 µm, (C) 10 µm. (D) Summary of the distribution of RhoGEF and RhoGAP proteins over different subcellular compartments.

## Focal adhesions are major sites of Rho GTPase regulation

To enhance the ability to resolve RhoGEF/RhoGAP localization to focal adhesions, a key site of cytoskeletal dynamics, we used total internal reflection fluorescence (TIRF) microscopy in COS-7 cells. This led to the identification of an unexpected one fourth (37) of all RhoGEFs and RhoGAPs that clearly associated with these structures, as determined by colocalization with the adhesion marker Paxillin. Efforts to define the components of integrin adhesions have already revealed a prevalence of Rho signaling network proteins(*23, 24*). 9 RhoGEFs/RhoGAPs were previously proposed to be an integral part of the adhesome (adhesome.org), of which we confirmed six. For 24 proteins identified in our screen no evidence for focal adhesion localization existed in literature (Table S3), among them well-studied regulators such as SOS, TRIO, CHN2 and KALRN (Fig. 3B, Fig. S4). Our data thus indicates that intricately tuned Rho signaling is more crucial in the control of cell-matrix adhesion than previously assumed.

Notably, six regulators exhibited a distinct microlocalization with a pericentric enrichment juxtapositioned to focal adhesions and a fluorescence intensity minimum in the center (Fig. S5). We also found that RhoGEFs and RhoGAPs with activity towards Rac1 were overrepresented at focal adhesions compared to RhoA and Cdc42 (Fig. S6, Tables S2, S3).

## Prevalence of actin-associated RhoGEFs and RhoGAPs

Next, we focused on identifying regulators associating with actin. We treated cells with Cytochalasin D, an agent that disrupts the actin network and induces the appearance of phalloidin-reactive filamentous foci (*25*). By scoring transiently expressed RhoGEFs and RhoGAPs for their colocalization with these foci, we identified 34 actin-associated proteins, including 23 regulators that have previously not been reported to bind actin, such as DOCK1, ITSN2 and ARHGAP5 (Fig. 3C and Fig. S7). This finding emphasizes the importance of close proximity of Rho regulators to actin to locally sense and control cytoskeletal dynamics. RhoGEFs and RhoGAPs with activity towards Cdc42 were overrepresented for association with actin relative to RhoA and Rac1 (Fig. S6).

Overall, we found 70% (99/141) of the overexpressed RhoGEF/RhoGAP proteins enriched at distinct subcellular compartments (Fig. 3D, Table S3). The vast majority of regulators thereby localize to structures previously shown to harbor Rho signaling. Notably, additional ‘non-canonical’ locations, comprising the Golgi, mitochondria, lysosomes, endomembranes and the endoplasmic reticulum, were only decorated by RhoGAPs. We thus did not find evidence for Rho activation at these structures. Compared to a control set of proteins from the Cell Atlas study (*26*), significantly more proteins in our screen localized to focal adhesions, actin and the plasma membrane (p-values < 2.2e-16, < 2.2e-16, 4.154e-07 and odds ratios=30.2, 14.5 and 2.9 respectively). Compared to a study that assessed localization in a similar manner as our study(*27*), the RhoGEFs/RhoGAPs in our set were significantly enriched in terms of discrete localization (i.e. not cytoplasm or nucleus; p-value=4.083e-09, odds ratio=3.0). For 27 (19%) of the regulators in our collection, there were no previous reports describing their localization, and 28 (20%) and 53 (37.5%) were not annotated in the Cell Atlas and Uniprot databases, respectively. For 38 proteins (27%) the localizations we found were different from those reported (Table S3). Together, our data provides a cellular heat-map of Rho regulation and establishes the critical role of the RhoGEFs/RhoGAPs in conveying spatial context to Rho signaling.

Using high-content microscopy(*28*), we found only a limited relationship between the substrate specificities and overexpression phenotypes of the RhoGEFs/RhoGAPs (Supplementary Text). Their expression thus results in a continuum of cell morphologies that contrasts the clear-cut phenotypes observed upon global expression of activated forms of the Rho GTPases(*29*), reflecting the promiscuity and differential subcellular distribution of the RhoGEFs and RhoGAPs.

## Interactome analysis reveals that many RhoGEFs/RhoGAPs associate in complexes

Next, we used affinity purification coupled to mass spectrometry to obtain a comprehensive network of 1292 interactions (silver set; 1082 novel) among 863 unique proteins (Fig. 4A, Table S4). Our dataset explores a so far largely uncharted part of the human interactome and connects Rho GTPases to several signaling pathways(*30*) and functional complexes involved in linking the actin cytoskeleton to critical cell functions (Fig. 4A, Fig. S8). Notably, in addition to 20 interactions between RhoGEFs or RhoGAPs and Rho effector proteins and 24 interactions with small GTPases, our network includes 66 unique interactions between RhoGEFs/RhoGAPs themselves, indicating a previously unrecognized extensive crosstalk among the Rho regulators (Fig. 4B, enrichment p-value for all <<0.001; Fig. S9). Both homo- and heterotypic interactions occurred, with fewer complexes between RhoGAPs (24 GEF/GAP, 28 GEF/GEF, 11 GAP/GAP and 3 GEF-ABR or GEF-BCR). Connections include complexes that localize to the same cellular compartment, such as FARP2-VAV1 on focal adhesions or ARHGAP28-SYDE1 on actin filaments, where they may act in concert to fine-tune Rho responses (Fig. 4B).

**Figure 4.**
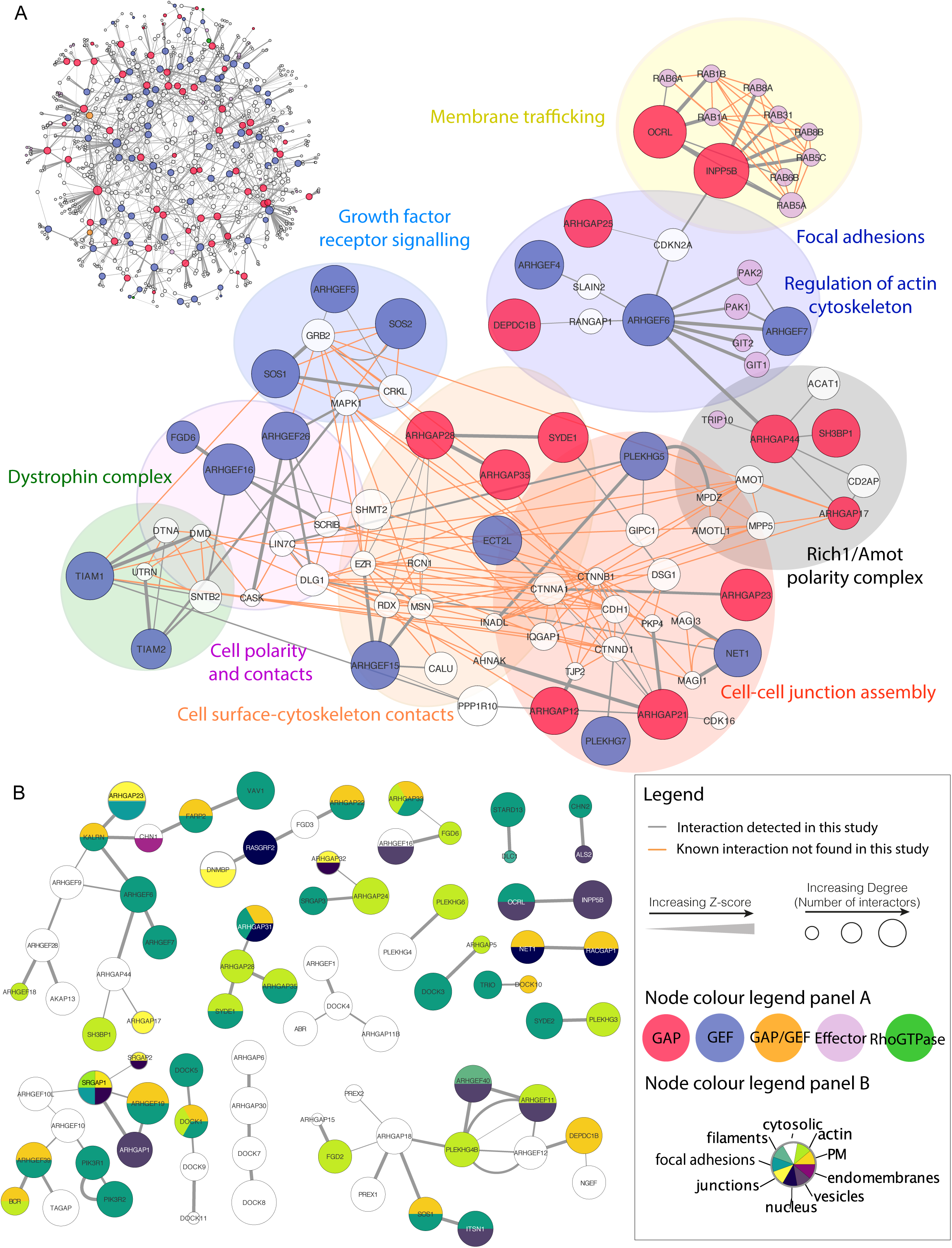
The RhoGEF/RhoGAP interactome is highly interconnected and includes components of multiple cellular processes. (A) RhoGEF/RhoGAP interactome network. As examples of subnetworks of the RhoGEF/RhoGAP interactome, interactions with complexes involved in cell polarity, junctions, membrane trafficking, growth factor receptor signaling and actin cytoskeleton organization are shown. (B) Bait-bait interactome. Our network is enriched (p-value <<0.001) in bait-bait interactions among RhoGEFs and RhoGAPs. Shown are all 66 interactions, with nodes color-coded by their subcellular localization (see also Figure 3).

**Figure 5.**
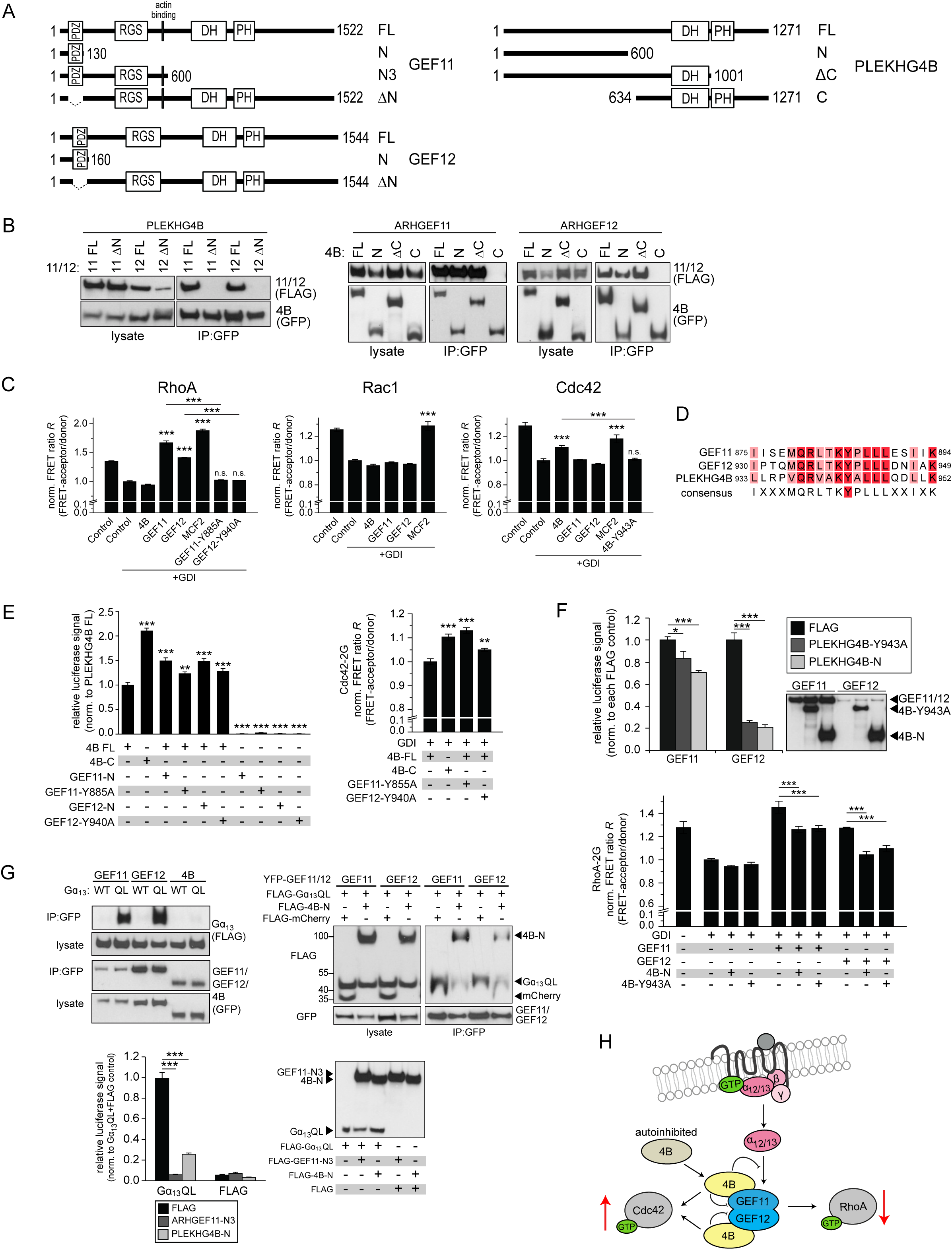
A multi-RhoGEF complex downstream of GPCR. (A) Schematic of the three human RhoGEFs and truncations used. (B) PLEKHG4B, ARHGEF11 and ARHGEF12 interact via their N-termini. Immunoprecipitations of indicated YFP-PLEKHG4B constructs with indicated FLAG-ARHGEF11 or FLAG-ARHGEF12 constructs from lysates of transfected HEK293T cells. (C) ARHGEF11 and ARHGEF12 are RhoA-specific GEFs, whereas PLEKHG4B activates Cdc42. FRET assay as described in Figure 1A and Figure S2. GEF-dead mutants: ARHGEF11-Y885A ARHGEF12-Y940A, PLEKHG4B-Y943A. (D) Sequence alignment of human ARHGEF11, ARHGEF12 and PLEKHG4B revealing the conserved tyrosine in the DH domain critical for catalysis(*50*). (E) Autoinhibition of PLEKHG4B and its release by ARHGEF11 and ARHGEF12 binding. Left panel: PLEKHG4B-stimulated SRE-luciferase reporter activation assay. Right panel: Cdc42-2G FRET ratios of HEK293T cells transfected with RhoGDI, mCherry-tagged PLEKHG4B and the indicated miRFP-tagged ARHGEF11 and ARHGEF12 mutants. (F) ARHGEF11 and ARHGEF12 are inhibited by PLEKHG4B. Upper panel: ARHGEF11/12-stimulated SRE-luciferase reporter activation assay. Anti-FLAG Western Blot shows the expression of the transfected constructs. Lower panel: RhoA-2G FRET ratios of HEK293T cells transfected with RhoGDI, mCherry-ARHGEF11/12 and the indicated miRFP-tagged PLEKHG4B mutants. (G) PLEKHG4B inhibits Gα13-mediated ARHGEF11/12 recruitment and downstream RhoA activation. Upper left panel: Immunoprecipitations of YFP-fusions of the three GEFs with wild type (WT) or constitutively active FLAG-Gα13 (QL) from lysates of transfected HEK293T cells. Upper right panel: Immunoprecipitations of YFP-tagged ARHGEF11/12 with indicated FLAG-tagged proteins from lysates of transfected HEK293T cells. Lower left panel: Gα13QL-stimulated SRE-luciferase reporter activation assay. Anti-FLAG Western Blot shows the expression of transfected constructs. (H) Model of PLEKHG4B interaction with ARHGEF11 and ARHGEF12 and their mutual regulation of Gα12/13-mediated GPCR signaling. All bar graphs show mean ± SD (n=3: C, E left, F, and G; n=4: E right). Significance was calculated by unpaired Student’s t-test (C, E right, F, and G) versus control+RhoGDI (C), versus PLEKHG4B (E right) or as indicated by lines. Significance was determined using One-way ANOVA, followed by Tukey’s multiple comparisons (E left). Significance was ranked as * p<0.05, ** p<0.01, *** p<0.001.

## Protein complexes controlling RhoGEF/RhoGAP localization

We found that the interactomes of focal adhesion-associated RhoGEFs/RhoGAPs are enriched in components of the adhesome(*24*) (Table S4, p-value=2.5e-08) and asked whether we could identify interactors that are responsible for their local recruitment. We selected ARHGAP39, a neuronal RacGAP controlling synaptic plasticity(*31*), and confirmed its interaction with PEAK1, a kinase implicated in cancer progression. PEAK1 also localized to focal complexes, as previously reported(*32*). The interaction critically involves a WW domain of ARHGAP39: when this domain is disrupted by a Y81A point mutation, it not only fails to associate with PEAK1 but is also no longer recruited to focal adhesions(Fig. S10). Likewise, the interactome of actin-localized regulators is enriched in actin-binding proteins (p-value=1e-13), whose function may be to similarly recruit these RhoGEFs/RhoGAPs (Table S4, Supplementary Text). C-DOCK RhoGEF subfamily regulators (DOCK6-8) were highly interconnected with the four members of the LRCH protein family (Fig. S11, Table S4). Indeed, we found a striking relocalization of the three RhoGEFs from the cytosol to the sites of LRCH expression: peripheral actin filaments and the endoplasmic reticulum. Given that RhoGEFs/RhoGAPs and their interactors are significantly enriched for PDZ, SH2, WW and SH3 domains (p-values <<0.01), which are common in scaffold or adaptor proteins, such recruitment likely affects the cellular distribution also of other Rho regulators.

## A multi-RhoGEF complex downstream of GPCR signaling mediates Rho GTPase crosstalk

To further study the interplay between Rho regulators, we functionally characterized a multi-RhoGEF complex identified in our proteomics analysis: the interaction between the yet undescribed RhoGEF PLEKHG4B and ARHGEF11 and ARHGEF12, two well-studied activators of RhoA signaling downstream of G protein-coupled receptors (GPCR) that have been shown to be essential for chemokine signaling-driven tumor cell invasion(*33, 34*) (Fig. 4B, Fig. 5A, Fig. S12A). ARHGEF11/12 are engaged at the membrane upon GPCR stimulation when activated heterotrimeric Gα12 and Gα13 subunits bind to the RGS (regulator of G protein signaling) domains of the GEFs(*35*). PLEKHG4B is one of the eight members of the Pleckstrin homology domain containing family G (PLEKHG) proteins. The N-termini of the three proteins are critical to this interaction (Fig. 5B). While ARHGEF11 and ARHGEF12 are RhoA-specific, PLEKHG4B selectively activated Cdc42 (Fig. 5C) and thus draws an additional Rho GTPase substrate into this multi-GEF assembly. We observed that PLEKHG4B is another RhoGEF that is subject to autoinhibition, as the truncation of the N-terminus of PLEKHG4B increased its Cdc42 GEF activity compared to the full-length form (Fig. 5E, Fig. S12B). An increase in the Cdc42 GEF activity of full-length PLEKHG4B appeared when we added either N-terminal fragments of ARHGEF11 or ARHGEF12 or catalytically inactive mutants (Fig. 5C-E). This suggests that binding to its partner GEFs releases the autoinhibition of PLEKHG4B. Conversely, coexpressing an N-terminal fragment of PLEKHG4B or its GEF-inactive mutant strongly decreased the catalytic activities of ARHGEF11 and ARHGEF12 towards RhoA (Fig. 5D,F). The three proteins thus mutually control one another’s GEF activities. In the context of the GPCR signaling pathway, both ARHGEF11 and ARHGEF12 selectively associated with constitutively active Gα13QL, as expected. Addition of PLEKHG4B, however, inhibited this interaction (Fig. 5G, upper panels). Fittingly, while the expression of Gα13QL robustly activated downstream Rho signaling through engagement of the endogenous RGS domain-containing RhoGEFs, this response was inhibited by the dominant negative ARHGEF11 N-terminus that competes with the endogenous GEFs for Gα13 binding(*36*). Importantly, the N-terminus of PLEKHG4B, also capable of binding to ARHGEF11/12, caused an equally strong inhibition of Gα13 signaling.

To our knowledge, this is the first description of a multi-Rho regulator assembly, consisting of exchange factors with different GTPase specificities. In complex, ARHGEF11 and ARHGEF12 enhance the Cdc42 GEF activity of PLEKHG4B. In return, PLEKHG4B inhibits ARHGEF11/12-mediated RhoA activation in two ways: directly by reducing their catalytic activity, and indirectly by perturbing their engagement with Gα12/13. We thus provide a proof of concept for a novel mechanism of cross-talk among Rho GTPases in a pathway implicated in cancer metastasis, that was previously thought to only activate RhoA(*37*) (Fig. 5H).

## Discussion

Rho GTPases orchestrate complex morphodynamic cell behaviors. A fundamental question is how the underlying highly localized processes are spatially maintained while the components of the Rho signaling system are subject to entropic leakage. Recent studies have challenged the perception of Rho signaling as a static, linear and ‘GTPase-centric’ process, in which a membrane-bound GTPase is sequentially regulated by a GEF and a GAP to control one specific cytoskeletal structure(*12, 13, 38, 39*). Here, we provide the first systems-level study of RhoGEF/RhoGAP function that establishes the framework for a dynamic ‘regulator-centric’ concept. A critical function of the RhoGEFs and RhoGAPs is to contextualize and spatially delimit the Rho signaling networks. They do so by providing positional cues based on the placement of the enzymes on dedicated supramolecular structures and the assembly of additional signaling proteins. Our data establishes focal adhesions and actin filaments as additional major sites of Rho signaling regulation.

We found that many regulators associate to collaborative networks, such as the PLEKHG4B/ARHGEF11/ARHGEF12 complex characterized here. This cooperativity of the RhoGEFs/RhoGAPs, together with their promiscuity towards their substrates, increases their combinatorial possibilities to engage multiple Rho family members simultaneously, and to fine-tune their activities. Most regulators are autoinhibited, presumably due to the back-folding of adjacent regions onto the catalytic domains, thereby preventing access to substrates(*40–42*). This suggests that the RhoGEFs/RhoGAPs are poised to respond to context-specific upstream stimuli and feedback regulation, a mechanism that can further spatially confine the activity of the regulators to micrometer-size zones while localizing on supramolecular structures. Diffusion spreads the activated GTPases from their source and thereby the local morphogenetic signals. This is balanced by an efficient turnover of the GDP/GTP cycle, mediated by the action of RhoGAPs in the vicinity of the activated GTPases. RhoGAPs are more promiscuous than RhoGEFs, a finding that is in agreement with previous biochemical studies on selected regulators(*43, 44*). They are also less interconnected in homotypic GAP/GAP complexes and have significantly fewer domains than RhoGEFs (p-value=5e-04), and are thus potentially more autonomous from regulation. Moreover, only RhoGAPs localize to ‘non-canonical’ structures that are not reported to host Rho signaling. These properties may contribute to a housekeeping function of the RhoGAPs, allowing them to efficiently reset the basal state of the GDP/GTP cycle and thus to prevent signals from leaking to the cell volume. This is reminiscent of other fundamental reaction cycles driven by the activities of opposing enzymes. Inactivating protein phosphatases, for example, also tend to display lower sequence selectivity than their kinase counterparts(*45*). RhoGDIs finally ensure the re-cycling of Rho GTPases through the confined activity zones.

The lack of Cdc42-GAP specificity is surprising. Cdc42 activity often peaks at early stages of cellular morphogenesis, for instance during cell polarization, protrusion or adhesion and may have a unique initiating role in these processes. A failsafe termination of the underlying signaling programs may require the rigorous simultaneous inactivation of the entire cascade of Rho GTPases involved, including Rac1 and RhoA.

The mechanisms governing local Rho signaling are fundamentally different from those in place in the Ras GTPase system, with only about 16 GEFs/GAPs(*46*). Here, spatial organization does not require the contribution of opposing regulators but instead arises from cycles of lipid anchoring and membrane release of the Ras proteins(*47–49*). The organization principles presented in this study thus add to the emerging landscape of mechanisms controlling localized activities of small GTPases.

This integrated study provides a basis by which the complexity of spatio-temporal control of Rho signaling can be further dissected and rules by which cells regulate context-specific functions can be explored.

## Supplemental Information

Supplemental Information includes 14 figures, 4 tables and 2 data files. This and extended information can be found at <u>Biostudies:S-BSST160</u> (*51*) and at http://the-rhome.com. The mass spectrometry proteomics data from the Q Exactive HF-X run have been deposited to the ProteomeXchange Consortium via the PRIDE (*52*) partner repository with the dataset identifier PXD010084. The rest of the mass spectrometry proteomics data have been deposited under the dataset identifier PXD010144.

## Author Contributions

Conceptualization: P.M.M., R.D.B., T.P., E.P., and O.R.; Methodology: P.M.M. (FRET and TIRF screen), O.R., C.W. (confocal screen), R.D.B., K.M.A., V.N., M.S., M.T., O. Popp and B.L. (Mass Spectrometry), J.R. (case study, Cytochalasin D screen), L.E.H., R.D.B., P.M.M., O.R., C. Bakal, E.P. (computational methods), S.Z., L.S., L.B., G.M., T.R., K.F., J.R. and P.T. (library); Analysis: E.P., P.M.M., J.R., R.D.B. and O.R.; Software: E.P., G.G. and P.M.M.; Investigation: R.L.E., L.B., M.T.C., C. Barth, R.W.W. and P.T.; Data Curation: E.P., P.M.M., R.D.B., J.R. and O.R.; Resources: K.C. and O. Pertz; Writing: O.R. with contributions of E.P. and P.M.M; Partial Supervision and Project Administration: E.P., K.C., P.M., F.P.R. and T.P.; Supervision: O.R.

## Acknowledgements

This work is dedicated to the memory of our dear mentor, Tony Pawson, without whom this study would not have been initiated. We thank O. Daumke, R. Hodge, D. Panakova and P. Bieling for critically reading the manuscript and R.D. Fritz for helpful discussions. We thank I. Laue, D. Heidler and the rest of the Rocks lab for technical assistance. We thank the MDC Advanced Light Microscopy Facility for technical assistance. This work was supported by the Human Frontier Science Program (LT000759/2008-L) (to O.R.), the CIHR Post-doctoral fellowship award (to R.D.B.), the Cancer Research UK (CRUK) Programme Foundation Award (C37275/A20146) and the Stand Up to Cancer campaign for Cancer Research UK (to C.B.), Genome Canada through Ontario Genomics, the Ontario Government (ORF GL2-025) and the Terry Fox Research Institute (to T.P). Interaction proteomics at the Lunenfeld-Tanenbaum Research Institute was performed at the Network Biology Collaborative Centre, currently supported by Genome Canada and Ontario Genomics (OGI-139).F.P.R. and E.P. were supported by the Canada Excellence Research Chairs Program, the Krembil Foundation, the Avon Foundation and by the NIH/NHGRI Center of Excellence in Genomic Science program (HG004233).

